# Inferring time-aware models of cancer progression using Timed Hazard Networks

**DOI:** 10.1101/2022.10.23.513436

**Authors:** Jian Chen

**Author notes:** Please address all correspondence to.

## Abstract

Analysis of the sequential accumulation of genetic events, such as mutations and copy number alterations, is key to understanding disease dynamics and may provide insights into the design of targeted therapies. Oncogenetic graphical models are computational methods that use genetic event profiles from cross-sectional genomic data to infer the statistical dependencies between events and thereby deduce their temporal order of occurrence. Existing research focuses mainly on the development of graph structure learning algorithms. However, no algorithm explicitly links the oncogenetic graph with the temporal differences of samples in an analytic way. In this paper, we propose a novel statistical framework Timed Hazard Networks (TimedHN), that treat progression times as hidden variables and jointly infers oncogenetic graph and pseudo-time order of samples. We modeled the accumulation process as a continuous-time Markov chain and developed an efficient gradient computation algorithm for the optimization. Experiment results using synthetic data showed that our method outperforms the state-of-the-art in graph reconstruction. We highlighted the differences between TimedHN and competing methods on a luminal breast cancer dataset and illustrated the potential utility of the proposed method. Implementation and data are available at https://github.com/puar-playground/TimedHN

## 1 Introduction

The genesis and progression of human cancer can be viewed as a Darwinian, multistep evolutionary process at the cellular level [1]. In this process, cancer cells acquire selective advantages by accumulating a series of genetic alterations (e.g., Somatic mutations, copy number alterations, changes in DNA methylation, gene expression, and protein concentration), resulting in clonal expansions [2]. The accumulations of genetic alterations in different patients often present a common pattern, for example, the sequential accumulation of APC→K-RAS→TP53 gene mutations in colorectal carcinogenesis [3]. However, the inference of more complicated dependencies within a larger number of genetic alterations is still an open question and could potentially impact patient treatment.

During the past 20 years, a dozen oncogenetic modeling methods have been developed for cross-sectional samples. Assuming different individuals’ genetic alteration profiles are independent observations from the same multivariate stochastic process, these methods construct directed graphical models that reflect the dependencies or causalities between genetic alterations among the patient population using cross-sectional samples. Specifically, each node stands for a genetic event whose probability depends on the events connected by incoming edges. Commonly used oncogenetic models infer three types of graphs. The first type of models infer a tree or forest structure where a single event may have multiple outgoing edges but have at most one incoming edge (e.g., oncotrees [4], METREX [5], Mtreemix [6], CAPRESE [7]). This assumption was made for modeling simplicity. Thus, these models only capture the dominant factors in oncogenesis and are expected to have a lower false-positive rate. The second model type infers a directed acyclic graph (DAG), where events may have multiple incoming edges without forming a cycle. (e.g., Conjunctive Bayesian Networks [8], DiProg [9], Bayesian Mutation Landscape [10], TO-DAG [11], CAPRI [12]). The third type of model uses a continuous-time Markov chain to infer a general directed graph without the acyclic constraint (e.g., NAM [13], Mutual Hazard Networks [14]). These methods have been used to model the accumulation of genetic alterations in many cancer types.

The main limitation of existing oncogenetic models is that they discard the progression times by either marginalization or heuristics to infer a static model. However, the temporal order represented by the progression times is informative for oncogenetic analysis. This paper proposes a novel statistical model called TimedHN that can jointly infer progression times and the oncogenetic graph. Specifically, following the Mutual Hazard Networks, the progression is modeled as a continuous-time Markov chain parameterized by a hazard network [14]. However, instead of marginalizing out the progression times, we included times as hidden variables in our objective function. The hazard network and progression times of samples are estimated by solving a constrained maximum likelihood problem using the backpropagation algorithm [15]. In addition, since the mutation profile has a long tail nature [16, 17], we developed an efficient method that can take advantage of data sparsity to compute the likelihood and its gradient in a subspace of all states, significantly reducing the model’s memory and time complexity. After the optimization, TimedHN can use the estimated hazard network to compute the maximum likelihood transition path and the expectation of progression time for each sample. In other words, it can estimate not only the temporal order of events but also the temporal order of samples.

The performance of the proposed method in terms of precision, recall, and F-score was compared to three state-of-art methods and the classic oncotrees in our benchmark experiments using synthetic data. In the simulations, we also compared TimedHN with itself, which uses real sampling times as constants to demonstrate the correctness and advantage of the proposed joint inference algorithm. Then, we conducted experiments using noisy data. The results demonstrated the robustness of our model to observation errors. Furthermore, the results of two simulation experiments showed that the time cost of our gradient computation algorithm is linear to the total number of events *n* and is exponential to the number of accumulated events *k*, which is usually much smaller than *n*. Finally, we applied our method on a luminal breast cancer dataset [18] to further demonstrate the performance of TimedHN.

## 2 Method

We model the event accumulation process as a continuous-time Markov chain parameterized by a weighted directed graph. The graph and progression times are inferred by constrained maximization of the log-likelihood using the backpropagation algorithm. In order to use cross-sectional data, our model assumed samples to be independent. Then, we developed an efficient algorithm that can compute the gradient without creating the full transition matrix for the exponential number of states, thus significantly reducing the computational complexity.

### 2.1 Model Overview

Following the Mutual Hazard Networks [14], we model the mutation accumulation of *n* genetic events in cancer progression as a continuous-time Markov chain (CTMC) on 2^*n*^ states. States are represented by *n* dimensional binary vectors **x** ∈ {0, 1}^*n*^, where **x**_*i*_ = 1 means that event *i* has occurred in the tumour by time *t*, while **x**_*i*_ = 0 means that it has not. We assumed that every progression trajectory starts at a normal state **x** = (0, 0, …, 0), accumulates *irreversible* genetic alteration events *one at a time*, and will eventually ends at a fully aberrant state **x** = (1, 1, …, 1). Observed sample profiles correspond to states at unknown intermediate times 0 *< t <* ∞ from independent progression.

Fig. 1 provided an overview of the method. Transition rates are parameterized by a hazard rate matrix **R**, which could be interpreted as an inter-event graph and spontaneous rates. Due to the accumulation assumptions, the transition rate matrix **Q** is constrained to be a hypercube. The likelihood of a sample at a given time *t* is just one entry on the first row of the transition probability matrix **P**(*t*), which is computed by the matrix exponential of the transition rate matrix **Q**.

**Figure 1:**
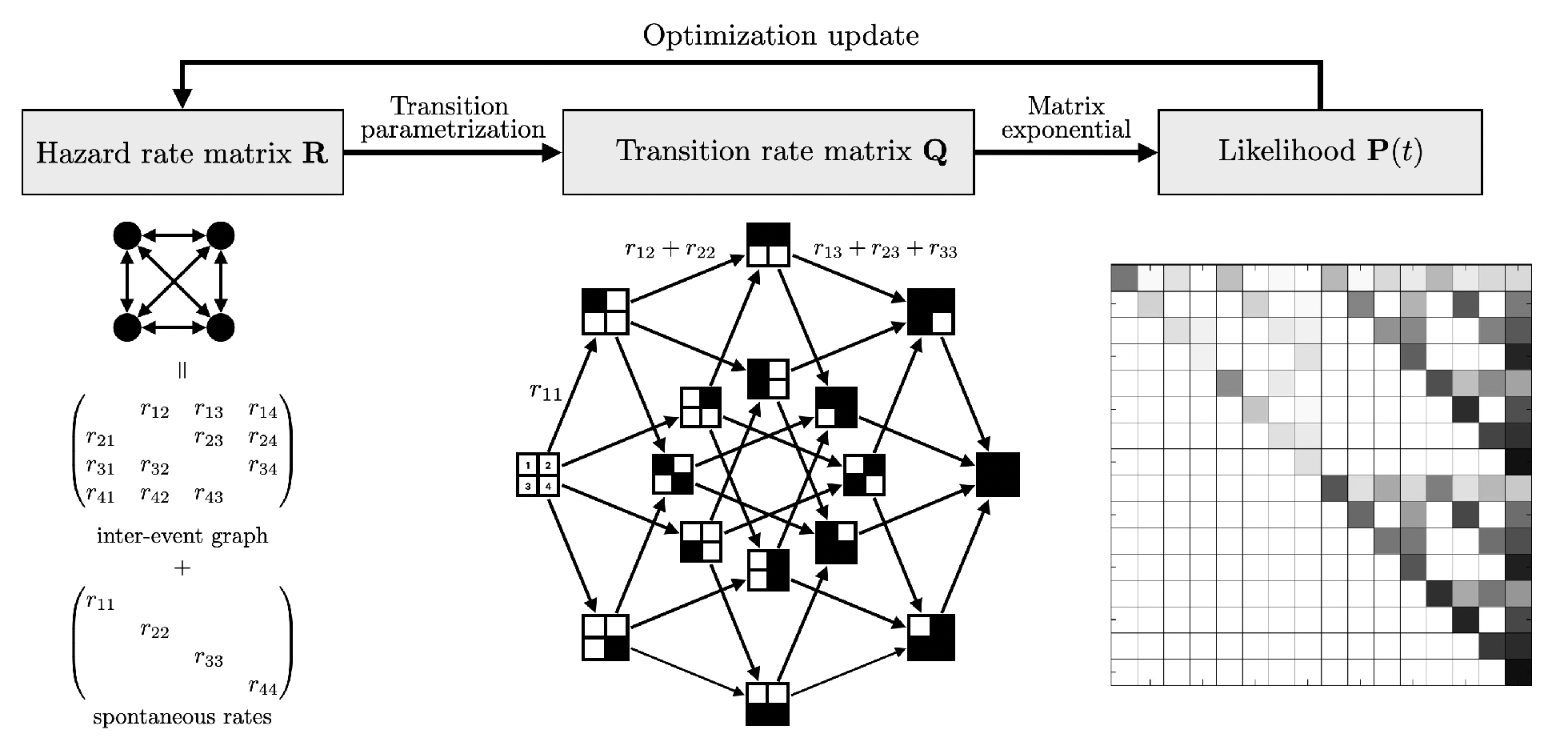
Overview of the proposed method

#### 2.1.1 Hazard network

We use a weighted directed graph (hazard network) with adjacency matrix **R** ∈ ℝ^+*n×n*^ to represent the pairwise dependencies and use the weights to parameterize the transition rate matrix describing the accumulation process. Specifically, the model was build with three assumptions: First, for any event *j*, its waiting time without being affected by other events has an exponential distribution *t*_*i*_|**0** ∼ Exp(*R*_*ii*_). We call it a spontaneous accumulation and *R*_*ii*_ is the spontaneous rate. Second, without considering the spontaneous accumulation, the waiting time of event *j* under the influence of event *i* also has an exponential distribution, *t*_*j*_|*i* ∼ Exp(*R*_*ij*_). Third, we assumed that the pairwise dependencies between events are independent of each other. Then, we can use these rates to model the conditional waiting time for any event, for example, for a state **x** = (…, *x*_*j−*1_, 0, *x*_*j*+1_, …) to acquire the event *j*, the conditional waiting time is the minimum of all the independnet waiting times: *t*_*j*_|**x** = min(*t*_*j*_|**0**, *t*_*j*_|*m*_1_, …, *t*_*j*_|*m*_*k*_), where *k* is the number of accumulated events and *m*_*j*_ is the index of the *j*-th happened event in state **x**. By the property of competing exponentials [19], *t*_*j*_|**x** is also exponentially distributed. Specifically, the distribution of the conditional waiting time is:

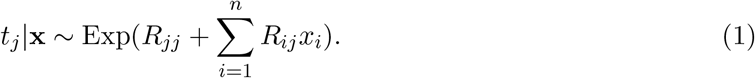

#### 2.1.2 Transition rate matrix

Next we show the event accumulation process is equivalent to a continuous-time Markov process on 2^*n*^ states that is uniquely defined by a transition rate matrix 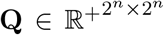. The states are ordered by a index function dec(**x**) = **x** · **b** + 1, where **b** = (2^0^, …, 2^*n−*1^)^*T*^ is the basis vector. Due to the progression assumptions: events are *irreversible* and accumulated *one at a time*, transition can only happen between two states that differ by one entry. For example, from state **x** = (…, *x*_*j−*1_, 0, *x*_*j*+1_, …) to state **x**_+*j*_ = (…, *x*_*j−*1_, 1, *x*_*j*+1_, …). The diagonal entries are defined as: 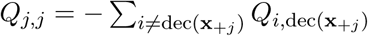 so that rows sum to zero, which is required for *Q* to be a valid transition rate matrix of a CTMC. The transition rate from state **x** to **x**_+*j*_ is defined as:

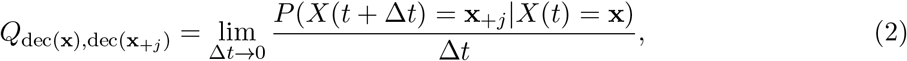

which by the definition of exponential distribution equals to the rate parameter of the waiting time in Eq. (1):

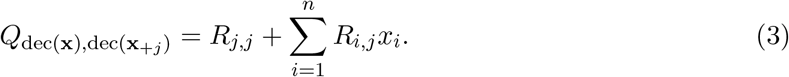

#### 2.1.3 Computation in subspace

The major problem of the computation in TimedHN is the exponentially increasing number of states, which results in unfeasible computational costs of the transition probability matrix **P**(**x**) = (*e*^*t***Q**^). We developed an efficient method that can take advantage of the sparsity of **x** to compute the likelihood and its gradient. Specifically, the transition matrix **Q** is transformed by a column permutation matrix **U** such that **U**^*⊤*^**QU** keeps the upper triangular structure. The dec(**x**)-th column is mapped to the column with the smallest possible column number. For a sample that accumulated *k* events, in the {*m*_1_, *m*_2_, …, *m*_*k*_} entries, the smallest possible column number is 2^*k*^. We can write the column permutation as 2 independent transpositions that swap the *i*-th column and the (bit_*k*_(*i* − 1) · **b**_sub_ + 1)-th column, for *i* = 1, …, 2^*k*^, where 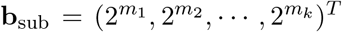 is the subspace basis vector and bit_*k*_(·) is the inverse function of dec(·) that map a integer to its *k* dimension binary vector. Thus, due to the upper triangular property [20, 21] (supplementary 1.1), we can get the likelihood by compute only the matrix exponential of the 2^*k*^-th order leading principal submatrix 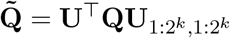 as:

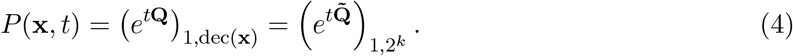

And the conditional time expectation is given by:

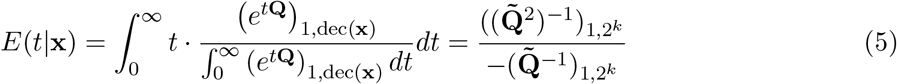

### 2.2 Optimization

We propose to jointly infer the hazard network that represents pairwise dependencies between events and progression times of samples through a constrained maximum likelihood estimation. The objective is given as:

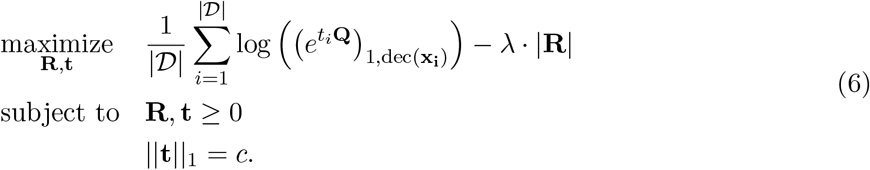

The likelihood of observing state **x** at time *t* is *P* (**x**, *t*) = (*e*^*t***Q**^)_1,dec(**x**)_, where the observation time *t* is considered as a hidden variable. Three constraints are added: First, the rates are non-negative, such that they are valid parameters of exponential distribution. Second, in order to prevent over fitting, we add *ℓ*1 regularization to encourage the sparseness of **R**, which leads to simpler topology of the hazard network. Third, observation times of all states are non-negative and have a constant summation. This constraint bound the times from above, which prevents the endless shrinkage of all hazard rates.

#### 2.2.1 Backpropagation in the subspace

We use the backpropagation algorithm to optimize the objective function. The partial derivative of the likelihood *P* (**x**, *t*) with respect to **R** and *t* are given as:

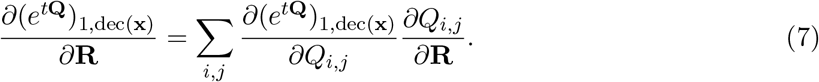

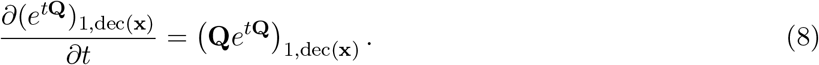

Since the likelihood equals to 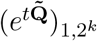 only depends on 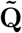, the derivatives could be computed as:

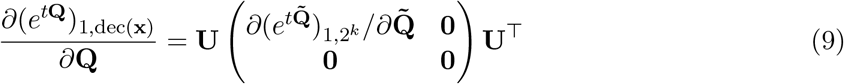

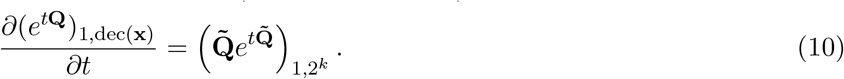

Finally, the derivative of matrix exponential is given as: 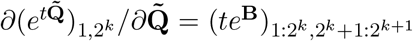 (supplementary 1.2 and 1.3), where

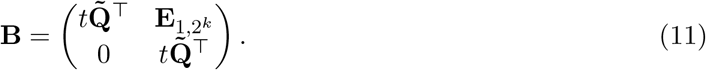

## 3 Experiment

### 3.1 Simulation on synthetic datasets

We sample synthetic data using CTMCs parameterized by random hazard networks to test the performance of TimedHN in inferring the structure of hazard networks using a given amount of data. We set the numbers of events to *n* = 15 and tested different sample sizes |D| ∈ {100, 250, 500, 1000}. We used hazard networks with forest and directed acrylic graph (DAG) structure to parameterize CTMCs. For each combination of sample size |D| and topology type, we generated 100 sets of data using different randomly generated hazard networks.

#### 3.1.1 Generation of random hazard networks

Forest are generated with a maximum depth of *log*(*n*). We randomly assign each node a depth between 1 to ⌊*log*(*n*)⌋, and make sure that each depth has at least one node. Finally, a parent is randomly selected from the nodes in the previous depth for each node. DAGs are generated with ⌊1.5*n*⌋ inter-event edges. Specifically, we assign topological sort ranks to the event nodes and randomly connect a higher-ranked node to a lower-ranked node to form edges. For modeling convenience, all edge weights are set to 1. The spontaneous rates of all the source nodes are set to 1, the spontaneous rates of all the rest nodes are set to 0.1.

#### 3.1.2 Event profiles sampling

In real-world data, we observed that most samples accumulate fewer than 10 mutations in cancerrelated genes. A possible explanation is that accumulating more mutations would make a cell less viable and thus rarely observed. We set the maximum number of accumulated events to 10 in the simulation based on this phenomenon. Specifically, we let a CTMC transition ten times to get one simulation run of the accumulation process. We then use the time *T* of the tenth jump as the termination time. Finally, we randomly sample an observation time *t* ∈ [0, *T*] and use the state of the CTMC at time *t* as a data sample. This sampling process is repeated multiple times to generate independent samples of synthetic datasets.

#### 3.1.3 Performance measure

Algorithmic performance was evaluated using the metrics precision, recall, and F-score between the inferred and true graphs used in the simulation. Precision and recall are defined as follows: precision tp/(tp+fp), recall tp/(tp + fn), and F-score 2tp/(2tp + fp + fn) which is the harmonic mean of precision and recall, where tp are the true positives, fp are the false positives and fn are the false negatives. Values for precision, recall, and F-score range from 0 to 1. The closer to 1, the better.

#### 3.1.4 Experimental settings

We compared our method with Mutual Hazard Networks (MHN) [14], [8], CAPRESE [7], CAPRI [12], and oncotrees [4]. For MHN, we used the source code downloaded from the paper websites. The L1 constraint weight for MHN is set to 1*/* |D| as suggested in the original paper. We used the implementation in the R package TRONCO (2.26.0) [22] with the default parameter settings for CAPRESE and CAPRI. For oncotrees, we used our python implementation. For TimedHN, we used the proposed method to maximize the average log-likelihood in Eq. (6) and set the learning rate to 1*e* − 3. We found that a larger regularization parameter *λ* usually results in more false negatives, which results in a low recall. On the other hand, although a smaller regularization parameter results in more false positives, the weights of true positive edges are usually much larger. Thus we can effectively remove false positive edges by using a threshold. In the simulation experiment, we used *λ* = 1*e* − 2, and we found 0.1 max(**R**) is a good choice for the threshold. However, the threshold is a hyperparameter and could be set manually after the optimization based on the user’s preference over precision and recall.

#### 3.1.5 Benchmark experiment on synthetic datasets

We compared the performance of TimedHN and four competing methods for inferring trees and DAGs using synthetic data. We also tested the TimedHN with true time observations instead if inferring times to demonstrate the advantage of the joint inference algorithm. Fig. 2 showed the performance of the six methods on simulations of 15 events with different sample sizes (|𝒟| ∈ {100, 250, 500, 1000}), obtained by averaging over 100 runs.

**Figure 2:**
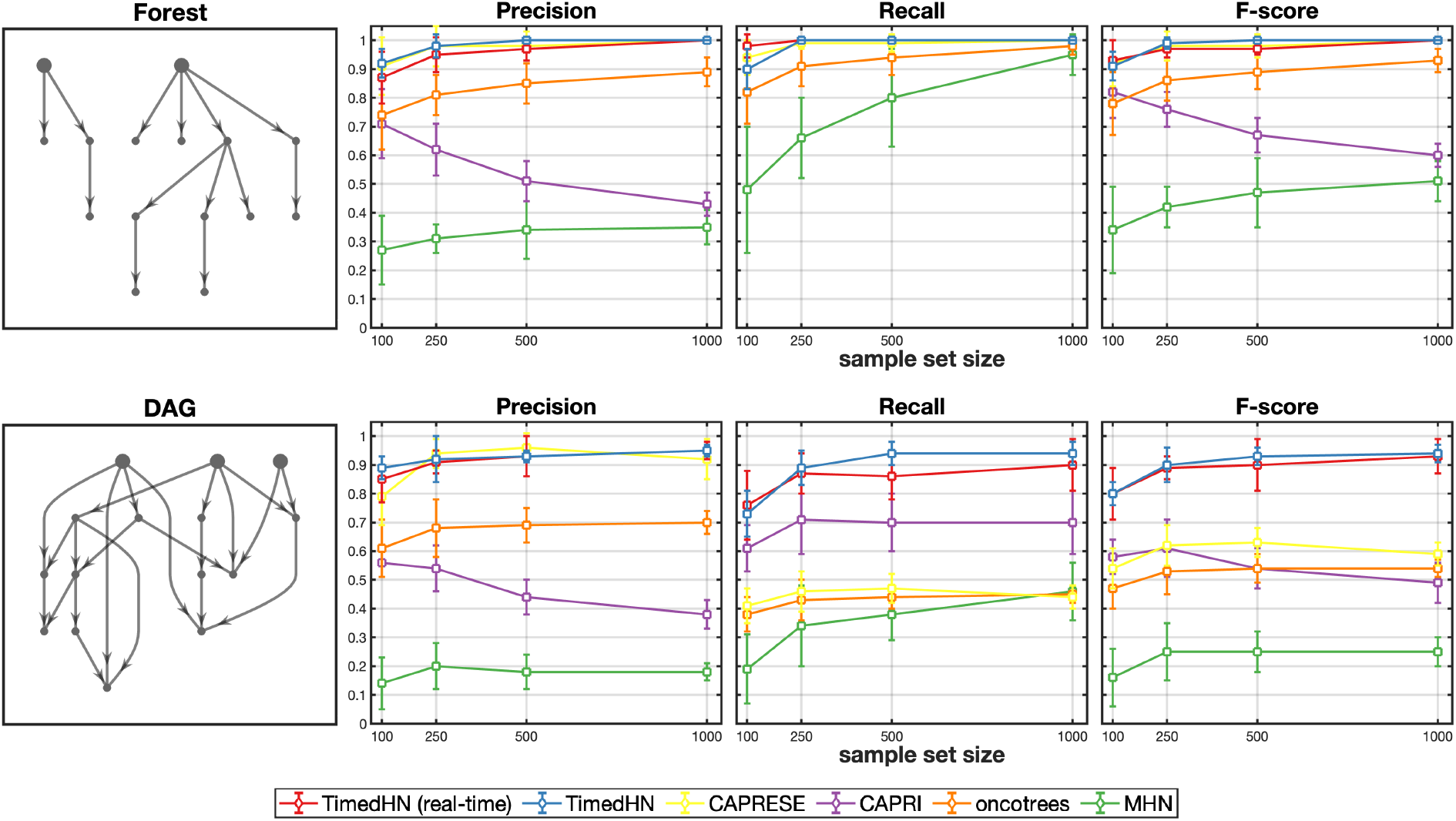
The precision, recall, and F-score of the six methods were compared using synthetic datasets consist of 15 events sampled from CTMCs parameterized by forests and DAGs.

We first applied the six methods to infer trees and forests, where every event has only one parent event. CAPRESE, and TimedHN using the real time and joint inference performed almost perfectly when sample sizes are larger than 500. Since CAPRESE was designed only to infer a tree or forest structure, this simulation perfectly fit its assumptions. Although oncotrees’ simple heuristic did not lead to perfect performance, it assumed a tree structure. Thus it still significantly outperformed CAPRI and MHN, which do not assume a tree structure. On the contrary, TimedHN managed to converge to the correct structure without these assumptions. We can see the precision and F-score of CAPRI decrease as the sample size increases. This is because CAPRI tends to infer a denser graph on larger sample sets, which results in a higher false positive rate and recall. We found MHN performed poorly even in this simple case. Two possible reasons are (i) it assumed identical distribution for progression times *P* (*t*) for all samples while the conditional distributions *P* (*t*|**x**) are different, which could lead to an erroneous topology of the hazard network. (ii) MHN also tries to infer negative hazard rates, which means the searching space of its optimization algorithm is much more complicated. Thus, it is easier to converge to a local optimum or result in overfitting. Moreover, in section 3.3, we found that MHN prefers to use edges with negative weights to fit the data rather than adding edges with positive weights.

Then, we applied the six methods to infer DAGs, where events can have multiple parent events. In terms of precision, CAPRESE is still comparable with TimedHN. However, in terms of recall, TimedHN using the real time or the joint inference outperformed all competing methods. TimedHN using the joint inference performed slightly better than using the true time and has a smaller standard deviation. A possible explanation is that the joint inference reduced the variance of sampling times. Due to the tree and forest assumption, oncotrees and CAPRESE could infer at most (*n* − 1) edges. Thus their recall is low when there are 1.5*n* edges in the true hazard networks. The recall of CAPRI decreased dramatically compared to their results on forests. Because its *prima facie causality* rules could fail in this case since the probability of observing a parent event is not guaranteed to be larger than that of observing a child. Finally, TimedHN significantly outperformed all competing methods in F-score due to its good performance on precision and recall.

#### 3.1.6 Experiment with profile noise

We performed simulation experiments to demonstrate the robustness of our model to observation errors and compared the results with the four competing methods. We used datasets of size |𝒟| = 250 generated by forest and DAG structures of *n* = 15 events. We randomly generated 100 datasets for each topology type using different hazard networks. Then we added noise by flipping each event independently with a small probability *ϵ* ∈ {0.1%, 0.25%, 0.5%, 1%, 2.5%, 5%}. Fig. 3 showed the performance of the six methods. Although the performance of all methods decreased as *ϵ* increased, TimedHN using real-time and joint inference are still comparable to CAPRESE in inferring forests and outperformed all the competing methods in inferring DAGs on every noise level. For forest structure, CAPRESE and oncotrees still outperformed CAPRI and MHN due to their structure assumptions. We found that the TimedHN had to infer some false edges to keep positive likelihoods for defected profiles. However, the weights of these false edges are usually small enough for a threshold to remove when the number of errors is small. If the amount of profile errors is large, the false positive edges would be indistinguishable from the edges connecting low-frequency events.

**Figure 3:**
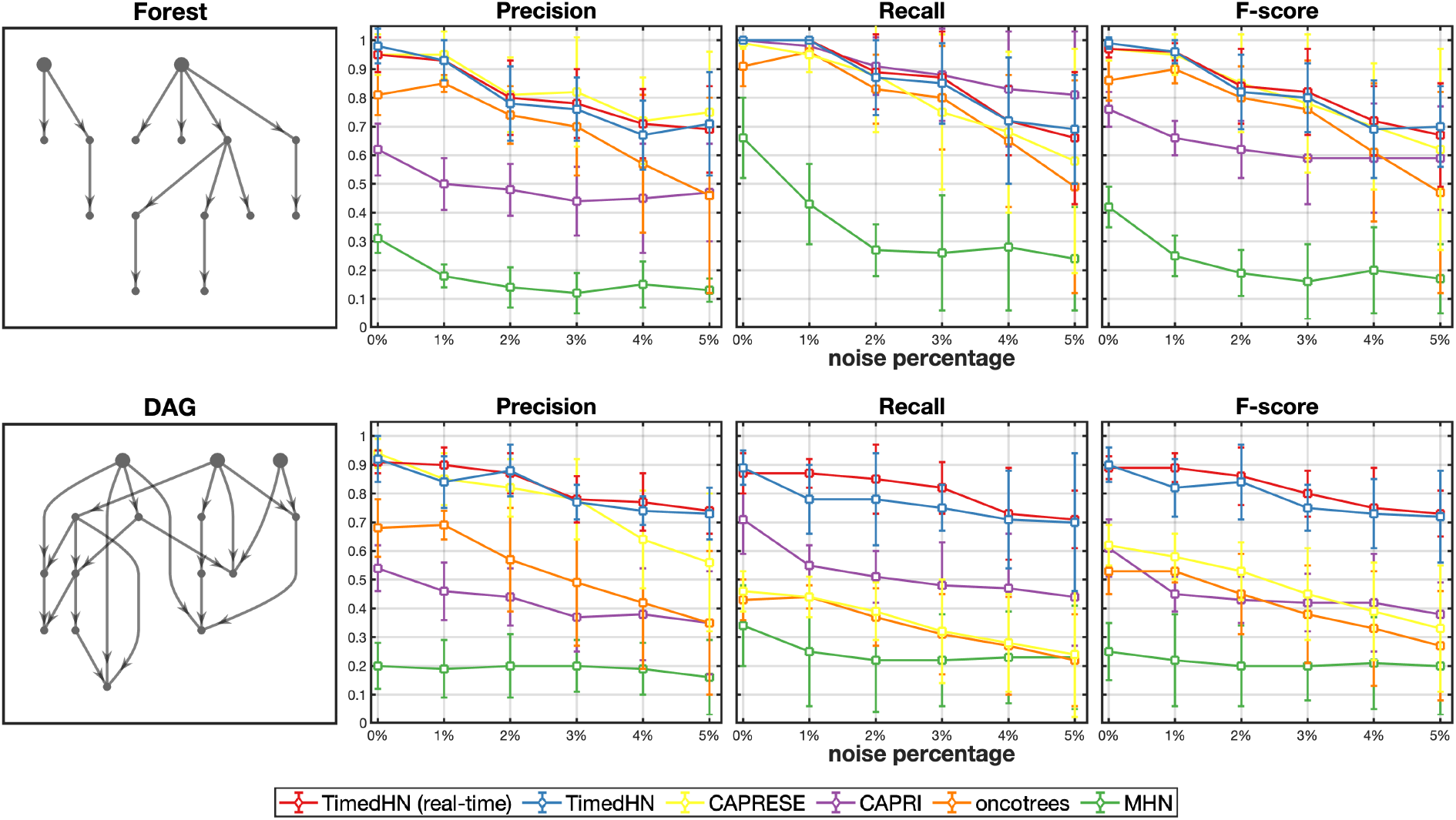
The precision, recall, and F-score of the six methods were compared using noisy synthetic datasets consist of 15 events sampled from CTMCs parameterized by forests and DAGs.

### 3.2 Time complexity analysis

We performed a test to analyze the time complexity of our gradient computation algorithm. We first fixed the profile dimension *n* = 20, and tested different numbers of accumulated events *k* ∈ {1, …, 15}. In Fig. 4(a), we can see the run time of gradient computation increased exponentially to *k*. It is because the computation of matrix exponential is the most time-consuming step, and the size of the matrix 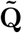 in section 2.2.1 grows exponentially to *k*.

**Figure 4:**
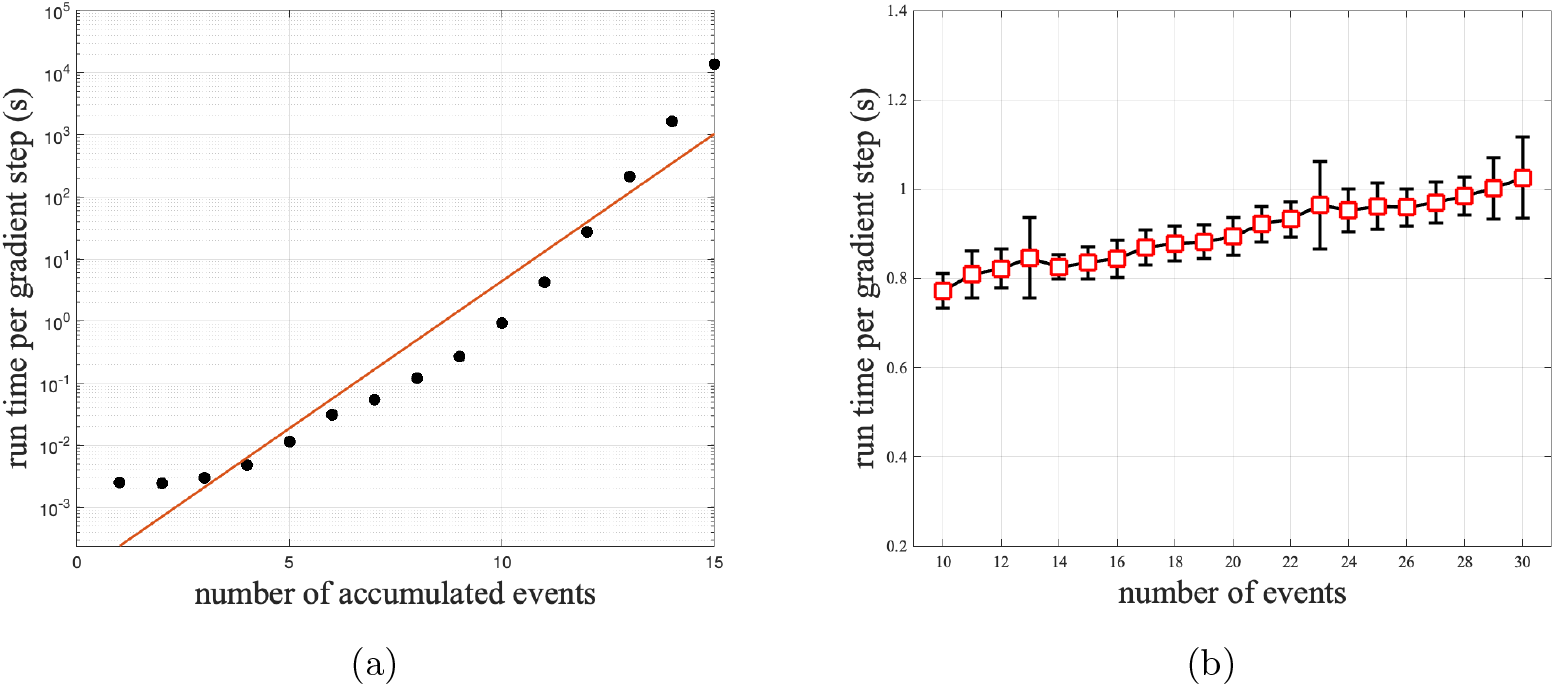
The time cos per gradient step is plot against different (a) number of accumulated events *k* and (b) number of events *n*.

Then we fixed the number of accumulated events *k* = 10, and tested different profile dimensions *n* ∈ {10, 11, …, 30}. In Fig. 4(b) we can see the computation time increased slowly as *n* increased. This is because since *k* is fixed, only the size of the permutation matrix **U** is increasing linearly to *n* using sparse representation. These results showed that our algorithm could take advantage of the sparsity of event profiles. When most patients just accumulated less than 10 cancer-related events, our algorithm can compute the gradient efficiently, regardless of the total number of events.

### 3.3 Luminal breast cancer

We applied TimedHN on luminal breast cancer [18] to further compare its performance with CAPRESE and MHN. The dataset was used in the CancerMapp pipeline [23] and is available at The Cancer Genome Atlas [24]. We extracted 685 profiles of luminal A and luminal B subtypes to form the dataset of this study. Events profiles were extracted from the Mutation Annotation Format (MAF) file used for the MutSig2CV [25] mutation analysis in The Cancer Genome Atlas, which cataloged mutations in 15889 genes in 973 breast tumor samples into five types of mutations: missense, Nonsense, in-frame indels, frameshift indels, and splice site. These mutations damage the function of a gene to different degrees by either changing the amino acids of the encoded protein or disable the translation process. Since its hard to evaluate the differences of this damages, we treat them equally as non-silent mutations of a gene. We used mutations of genes selected by the CancerMapp pipeline [23] for our experiments. It used a statistical approach to determine whether a gene mutation showed a significant change along the progression model inferred using expression profiles. Then, it used the Benjamini-Hochberg procedure [26] to compute a false discover rate (FDR) for each gene. In our experiment, for ease of interpretation, we only selected the genes that have an FDR lower than 0.01 (supplementary Table. 1).

We compared the results of TimedHN (Fig. 5) with the results of CAPRESE (Fig. 6(a)) since it showed the highest level of precision in the benchmark experiment. TimedHN captured all edges inferred by CAPRESE except PIK3CA→TP53. Instead of modeling the causality from PIK3CA, it inferred TP53 as a source node with a large spontaneous rate. This is to fit the 36 samples in the dataset with TP53 mutations but without PIK3CA mutations. The inferred independence of PIK3CA and TP53 are consistent with their different roles as oncogene and tumor suppressor, respectively, suggesting that abnormality in either gene can promote a malignant phenotype [27]. TimedHN also captured a few additional edges, such as: GATA3→CBFB, CBFB→GRHL2, CDH1→MED23, MAP3K1→CTCF, MAP2K4→ADAM29, MAP2K4→MYO6, etc. Although some of them are not as significant as edges also inferred by CAPRESE, these edges are strong enough to compensate the *ℓ*_1_ norm regularization and are necessary for the likelihood optimization of low frequency samples. Therefore, they also reflect non-negligible patterns of the dataset.

**Figure 5:**
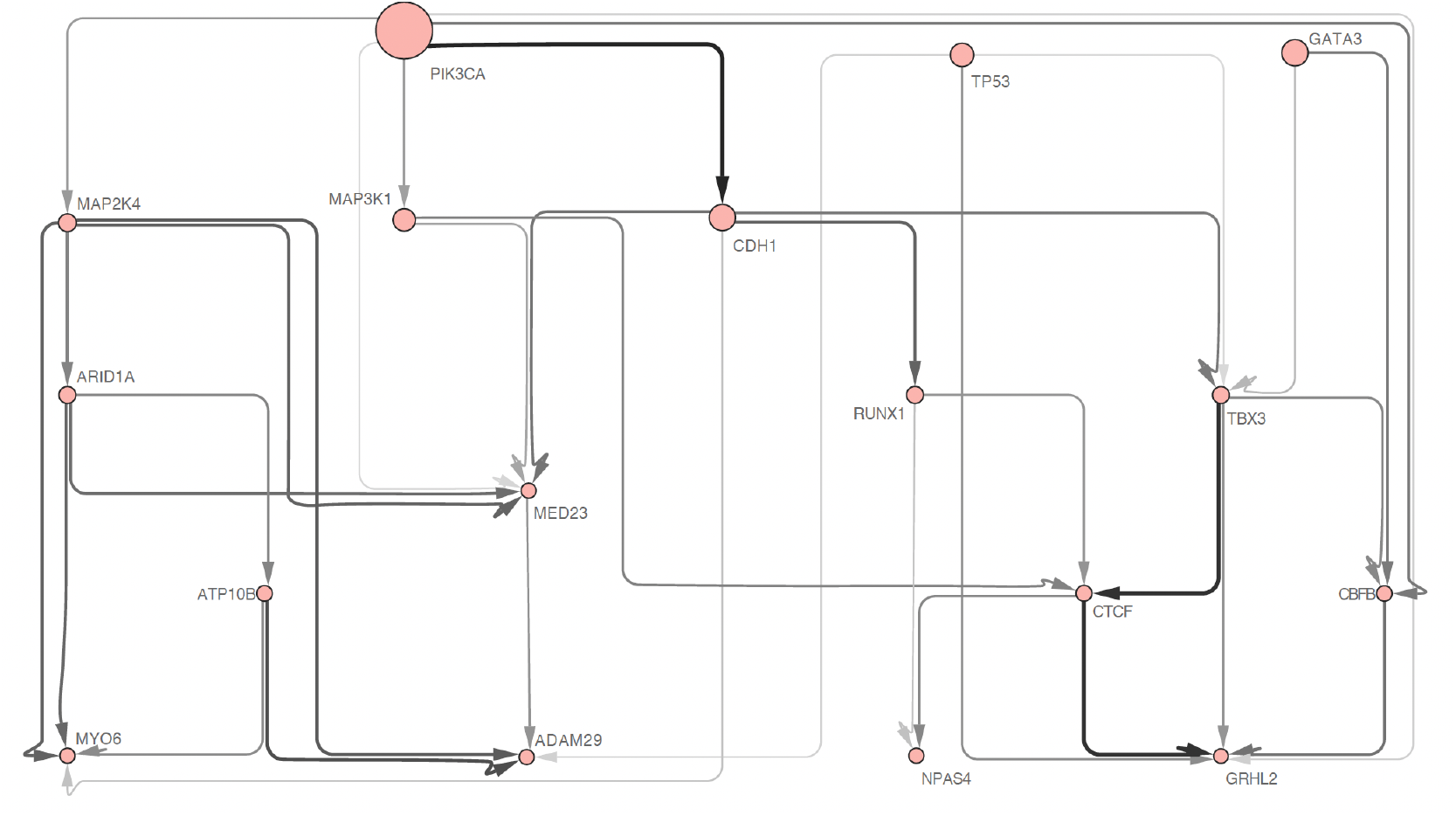
The oncogenetic graph inferred by TimedHN. Edge widths and shade are linear to the inter-event hazard rates. The node sizes are linear to spontaneous hazard rates.

**Figure 6:**
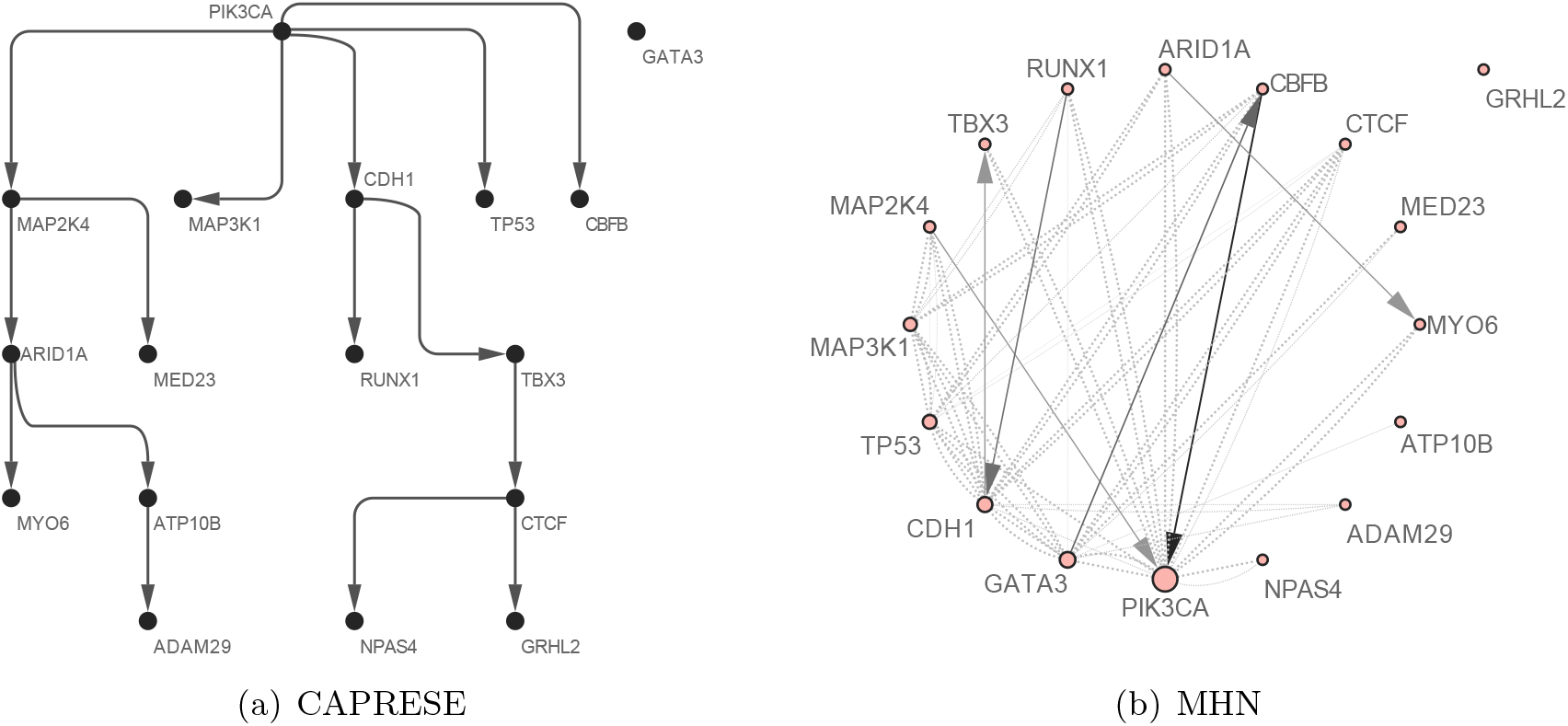
The oncogenetic tree inferred by CAPRESE is shown in subfigure (a) Its edges and nodes are shown in uniform width and size. The Oncogenetic graph inferred by MHN is shown in subfigure (b). Its edge widths and shade are linear to the exponential of inter-event hazard rates. Node sizes are linear to the exponential of spontaneous hazard rates. Solid lines represent edges with positive weights, and dashed lines represent negative ones.

We also compared with the results of Mutual Hazard Networks (Fig. 6(b)) to demonstrate its limitation. MHN inferred many mutually exclusive relationships using fully connected subgraph with only negative weighted edges, e.g., the mutual exclusiveness between MAP3K1, MAP2K4, and CDH1. However, it is expensive in terms of *ℓ*_1_ cost. As a result, the model tends to trade positive edges to model such a dense subgraph. Moreover, due to the incorrect assumption of the conditional time distribution *P* (*t*|**x**), the direction of some inferred edge with positive weight is reversed compared to the results from TimedHN and CAPRESE. On the other hand, methods like CAPRESE and TimedHN that only infer positive dependencies could also represent the mutually exclusive relationships by disconnection. Moreover, allowing negative hazard rates will significantly expand and complicate the searching space in the optimization, which could lead to the convergence to a local optimum or over-fitting.

Finally, in supplementary Fig. 1 demonstrated the ability of TimedHN to estimate pseudo-time orders of profiles and events. Event profiles were sorted by the conditional time expectation. A brutal force search of all possible accumulation orders of events in a profile was used to find the maximum likelihood accumulation order.

## 4 Conclusion

In this work, we have introduced a novel framework called TimedHN for the inference of the temporal order of samples and the oncogenetic graph (hazard network) underlying the accumulation of genetic events in cancer progression. Specifically, we included the progression time of the continuous-time Markov chain as a hidden variable in the optimization and developed an efficient algorithm to jointly infer the hazard network and the progression time through constrained optimization of the sample log-likelihood. The proposed optimization algorithm can take advantage of data sparsity and reduce the computational complexity from *O*(*e*^*n*^) to *O*(*e*^*k*^), where *n* is the total number of events and *k* is the number of accumulated events in a profile. We proved the correctness of TimedHN by showing convergence to the correct typology in simulation tests. Moreover, we compared TimedHN to the standard tree reconstruction algorithm based on co-occurrence patterns (i.e., Oncontree and CAPRESE) and a more general bayesian probabilistic graphical model (i.e., CAPRI) and the Mutual Hazard Networks (MHN). The results showed that TimedHN outperforms the state-of-the-art on synthetic data. Moreover, the advantage holds in the presence of mild observation noise in the event profiles. Furthermore, we experimented on luminal breast cancer mutation data using CAPRESE, MHN, and TimedHN. The analysis suggested that the results of TimedHN are highly consistent with the most precise method in the simulation (i.e., CAPRESE) and can infer novel dependencies that would remain undetected. At the same time, the analysis of the result of MHN showed its limitation in reliability and ease of interpretation.

TimedHN still has some limitations. Its application depends highly on event selection methods because it infers meaningful results only on a preselected set of genetic events that are supposed to be involved in a cumulative causal process. However, this is the limitation shared by all oncogenetic graph learning methods. TimedHN could serve as a tool to provide computational evidence for such hypotheses or to identify potential oncogenetic dependencies. Another limitation is the computational complexity of the proposed algorithm is still exponential to the number of accumulated events. Therefore, it is only efficient for sparse profiles.

Several future research directions are possible. First, the application of TimedHN is not limited to the analysis of cross-sectional data. It also applies to multi-region and single-cell data because the framework can compute the transition probability between any two states. For example, for a state **x** with *k* accumulated events, if an intermediate state with *k* − ∆*k* accumulated events is observed from the same patient, The computational cost in computing the likelihood of **x** will be reduced from *O*(*e*^*k*^) to *O*(*e*^*k−*∆*k*^ + *e*^∆*k*^). This property offers an opportunity for TimedHN to utilize datasets from different sources and benefit from the results of phylogenetic analysis [28, 29, 30].

Second, although the progression time of each sample is not observable, it can use the pseudotimes provided by established trajectory inference methods using expression profiles of single-cell data [31], and cross-sectional data [23]. TimedHN can treat pseudo-times as constant and only infer the oncogenetic graph. On the other hand, it can use pseudo-times to initialize the optimization. In this way, the proposed framework could be combined with trajectory inference methods as a multi-view learning [32] method that uses both mutation profiles and expression profiles for progression modeling.

Third, the proposed methods could be used for the analysis of other disease progressions, such as the development of drug resistance-associated mutations in the HIV genome [33] or similar cumulative causal processes.

In summary, this paper introduced a new, time-aware framework based on the continuous-time Markov chain that can utilize or infer the temporal information of the samples and the oncogenetic graph for modeling the accumulation of genetic events in cancer progression. We expect that in the future, TimedHN could be an essential building block in integrating phylogenetic and trajectory inference methods with oncogenetic models and aid the study of cancer.

## References

[1] Peter C Nowell. The clonal evolution of tumor cell populations: Acquired genetic lability permits stepwise selection of variant sublines and underlies tumor progression. Science, 194(4260):23–28, 1976.

[2] Mel Greaves and Carlo C Maley. Clonal evolution in cancer. Nature, 481(7381):306–313, 2012.

[3] Eric R Fearon and Bert Vogelstein. A genetic model for colorectal tumorigenesis. Cell, 61(5):759–767, 1990.

[4] Richard Desper, Feng Jiang, et al. Inferring tree models for oncogenesis from comparative genome hybridization data. Journal of Computational Biology, 6(1):37–51, 1999.

[5] Richard Desper, Javed Khan, et al. Tumor classification using phylogenetic methods on expression data. Journal of Theoretical Biology, 228(4):477–496, 2004.

[6] Niko Beerenwinkel, Jörg Rahnenführer, et al. Mtreemix: a software package for learning and using mixture models of mutagenetic trees. Bioinformatics, 21(9):2106–2107, 2005.

[7] Loes Olde Loohuis, Giulio Caravagna, et al. Inferring tree causal models of cancer progression with probability raising. PloS ONE, 9(10):e108358, 2014.

[8] Moritz Gerstung, Michael Baudis, et al. Quantifying cancer progression with conjunctive bayesian networks. Bioinformatics, 25(21):2809–2815, 2009.

[9] Hossein Shahrabi Farahani and Jens Lagergren. Learning oncogenetic networks by reducing to mixed integer linear programming. PloS ONE, 8(6):e65773, 2013.

[10] Navodit Misra, Ewa Szczurek, et al. Inferring the paths of somatic evolution in cancer. Bioinformatics, 30(17):2456–2463, 2014.

[11] Paola Lecca, Nicola Casiraghi, et al. Defining order and timing of mutations during cancer progression: the to-dag probabilistic graphical model. Frontiers in Genetics, 6:309, 2015.

[12] Daniele Ramazzotti, Giulio Caravagna, et al. Capri: efficient inference of cancer progression models from cross-sectional data. Bioinformatics, 31(18):3016–3026, 2015.

[13] Marcus Hjelm, Mattias Höglund, et al. New probabilistic network models and algorithms for oncogenesis. Journal of Computational Biology, 13(4):853–865, 2006.

[14] Rudolf Schill, Stefan Solbrig, et al. Modelling cancer progression using mutual hazard networks. Bioinformatics, 36(1):241–249, 2020.

[15] Robert Hecht-Nielsen. Theory of the backpropagation neural network. In Neural networks for perception, pages 65–93. Elsevier, 1992.

[16] Joshua Armenia, Stephanie AM Wankowicz, et al. The long tail of oncogenic drivers in prostate cancer. Nature genetics, 50(5):645–651, 2018.

[17] Hussein Mohsen, Vignesh Gunasekharan, et al. Network propagation-based prioritization of long tail genes in 17 cancer types. Genome Biology, 22(1):1–21, 2021.

[18] Michail Ignatiadis and Christos Sotiriou. Luminal breast cancer: from biology to treatment. Nature Reviews Clinical Oncology, 10(9):494–506, 2013.

[19] K Balakrishnan. Exponential distribution: theory, methods and applications. Routledge, 2019.

[20] Charles Van Loan. The sensitivity of the matrix exponential. SIAM Journal on Numerical Analysis, 14(6):971–981, 1977.

[21] Luca Dieci and Alessandra Papini. Padé approximation for the exponential of a block triangular matrix. Linear Algebra and its Applications, 308(1-3):183–202, 2000.

[22] Luca De Sano, Giulio Caravagna, et al. Tronco: an r package for the inference of cancer progression models from heterogeneous genomic data. Bioinformatics, 32(12):1911–1913, 2016.

[23] Yijun Sun, Jin Yao, et al. Computational approach for deriving cancer progression roadmaps from static sample data. Nucleic Acids Research, 45(9):e69–e69, 2017.

[24] DCFR Koboldt, Robert Fulton, et al. Comprehensive molecular portraits of human breast tumours. Nature, 490(7418):61–70, 2012.

[25] Michael S Lawrence, Petar Stojanov, et al. Discovery and saturation analysis of cancer genes across 21 tumour types. Nature, 505(7484):495–501, 2014.

[26] Yoav Benjamini and Yosef Hochberg. Controlling the false discovery rate: a practical and powerful approach to multiple testing. Journal of the Royal statistical society: series B (Methodological), 57(1):289–300, 1995.

[27] Bhuvanesh Singh and Pabbathi G others Reddy. p53 regulates cell survival by inhibiting pik3ca in squamous cell carcinomas. Genes & development, 16(8):984–993, 2002.

[28] Russell Schwartz and Alejandro A Schäffer. The evolution of tumour phylogenetics: principles and practice. Nature Reviews Genetics, 18(4):213–229, 2017.

[29] Sayaka Miura, Tracy Vu, et al. A phylogenetic approach to study the evolution of somatic mutational processes in cancer. Communications Biology, 5(1):1–11, 2022.

[30] Niko Beerenwinkel, Roland F Schwarz, et al. Cancer evolution: mathematical models and computational inference. Systematic Biology, 64(1):e1–e25, 2015.

[31] Wouter Saelens, Robrecht Cannoodt, Helena Todorov, and Yvan Saeys. A comparison of single-cell trajectory inference methods. Nature Biotechnology, 37(5):547–554, 2019.

[32] Chang Xu, Dacheng Tao, et al. A survey on multi-view learning. arXiv preprint 1304.5634, 2013.

[33] Niko Beerenwinkel, Martin Däumer, et al. Estimating hiv evolutionary pathways and the genetic barrier to drug resistance. The Journal of Infectious Diseases, 191(11):1953–1960, 2005.

