## supplemental file for "Inferring time-aware models of cancer progression using Timed Hazard Networks"

### TimedHN supplementary

Jian Chen

### 1 Model optimization

#### 1.1 Matrix exponential for block upper triangular matrix

If a square matrix $\mathbf{A}$ is block upper triangular [1, 2],

$$\mathbf{A} = \begin{pmatrix} \mathbf{A}_{11} & \mathbf{A}_{12} \\ 0 & \mathbf{A}_{22} \end{pmatrix}, \tag{1}$$

where $\mathbf{A}_{11}$ and $\mathbf{A}_{22}$ are square matrices. The matrix exponential has the same structure.

$$e^{\mathbf{A}} = \begin{pmatrix} e^{\mathbf{A}_{11}} & \mathbf{F} \\ 0 & e^{\mathbf{A}_{22}} \end{pmatrix}. \tag{2}$$

Where:

$$F = \int_0^1 e^{(1-u)\mathbf{A}_{11}} \mathbf{A}_{12} e^{u\mathbf{A}_{22}} du. \tag{3}$$

#### 1.2 Derivatives of matrix exponential

For an order $n$ square matrix $t\mathbf{A}$, the partial derivative of the $(i, j)$-th entry of its matrix exponential $e^A \in \mathbb{R}^{+n \times n}$ with respect to $\mathbf{A}$ is defined as:

$$\frac{\partial (e^{t\mathbf{A}})_{ij}}{\partial \mathbf{A}} = \begin{pmatrix} \partial(e^{t\mathbf{A}})_{ij}/\partial A_{11} & \cdots & \partial(e^{t\mathbf{A}})_{ij}/\partial A_{1n} \\ \vdots & \ddots & \vdots \\ \partial(e^{t\mathbf{A}})_{ij}/\partial A_{n1} & \cdots & \partial(e^{t\mathbf{A}})_{ij}/\partial A_{nn} \end{pmatrix}, \tag{4}$$

which is related to the first directional direvative (Gateaux derivative) of matrix exponential:

$$\frac{\partial (e^{t\mathbf{A}})}{\partial A_{ij}} = \lim_{h \to 0} \frac{1}{h} \left( e^{t(\mathbf{A}+h\mathbf{E}_{i,j})} - e^{t\mathbf{A}} \right) \tag{5}$$

$$= \int_0^t e^{(t-\tau)\mathbf{A}} \mathbf{E}_{i,j} e^{\tau \mathbf{A}} d\tau, \tag{6}$$

where $\mathbf{E}_{i,j}$ is a direction matrix, it has only one non-zero value at the $(i, j)$-th entry which equals to 1. According to the *proposition* 6.1 in [3],

$$\frac{\partial (e^{\mathbf{A}})_{ij}}{\partial \mathbf{A}} = \frac{\partial e^{\mathbf{A}^\top}}{\partial A_{ij}}. \tag{7}$$

Thus, we have the closed form solution for Eq.(4):

$$\frac{\partial (e^{t\mathbf{A}})_{i,j}}{\partial \mathbf{A}} = \int_0^t e^{(t-\tau)\mathbf{A}^\top} \mathbf{E}_{i,j} e^{\tau \mathbf{A}^\top} d\tau \tag{8}$$

##### 1.3 Efficient derivative computation

Since the numerical computation of the integral in Eq. (8) is time comsuming, we give an analytic solution for it.

$$\frac{\partial(e^{t\mathbf{A}})_{i,j}}{\partial\mathbf{A}} = \int_0^t e^{(t-\tau)\mathbf{A}^\top}\mathbf{E}_{i,j}e^{\tau\mathbf{A}^\top}d\tau \tag{9}$$

$$= \int_0^1 e^{(t-tu)\mathbf{A}^\top}\mathbf{E}_{i,j}e^{tu\mathbf{A}^\top}d(tu). \tag{10}$$

$$= t\int_0^1 e^{(1-u)(t\mathbf{A}^\top)}\mathbf{E}_{i,j}e^{u(t\mathbf{A}^\top)}du. \tag{11}$$

Following Eq. (3) and Eq. (11), we can then construct an order $2n$ square matrix $\mathbf{B}$:

$$\mathbf{B} = \begin{pmatrix} t\mathbf{A}^\top & \mathbf{E}_{i,j} \\ 0 & t\mathbf{A}^\top \end{pmatrix}, \tag{12}$$

And compute the derivative as:

$$te^{\mathbf{B}} = \begin{pmatrix} te^{t\mathbf{A}^\top} & \partial(e^{t\mathbf{A}})_{ij}/\partial\mathbf{A} \\ 0 & te^{t\mathbf{A}^\top} \end{pmatrix}. \tag{13}$$

#### 2 Luminal breast cancer

Table 1: Genes selected by CancerMapp pipeline that showed significant changes along the progression of gene expression profiles.

| Gene | Full name |
|---|---|
| PIK3CA | phosphoinositide-3-kinase, catalytic, alpha polypeptide |
| TP53 | tumor protein p53 |
| CDH1 | cadherin 1, type 1, E-cadherin (epithelial) |
| GATA3 | GATA binding protein 3 |
| MAP3K1 | mitogen-activated protein kinase kinase kinase 1 |
| MAP2K4 | mitogen-activated protein kinase kinase 4 |
| RUNX1 | runt-related transcription factor 1 (acute myeloid leukemia 1; aml1 oncogene) |
| TBX3 | T-box 3 (ulnar mammary syndrome) |
| CBFB | core-binding factor, beta subunit |
| CTCF | CCCTC-binding factor (zinc finger protein) |
| ADAM29 | ADAM metallopeptidase domain 29 |
| ARID1A | AT rich interactive domain 1A (SWI-like) |
| MYO6 | myosin VI |
| NPAS4 | neuronal PAS domain protein 4 |
| MED23 | mediator complex subunit 23 |
| ATP10B | ATPase, class V, type 10B |
| GRHL2 | grainyhead-like 2 (Drosophila) |

| Time expectation | Order | PIK3CA | TP53 | CDH1 | GATA3 | MAP3K1 | MAP2K4 | RUNX1 | TBX3 | CBFB | CTCF | ADAM29 | ARID1A | MYO6 | NPAS4 | MED23 | ATP10B | GRHL2 |
|---|---|---|---|---|---|---|---|---|---|---|---|---|---|---|---|---|---|---|
| 1.39536220437563 | R | | | | | | | | | | | | | | | | | |
| 2.59781465108044 | R --> MAP2K4 | | | | | |■| | | | | | | | | | | |
| 2.63675342915226 | R --> TBX3 | | | | | | | |■| | | | | | | | | |
| 2.64672276931439 | R --> ARID1A | | | | | | | | | | | |■| | | | | |
| 2.67017171718461 | R --> CTCF | | | | | | | | | |■| | | | | | | |
| 2.68515777276158 | R --> ATP10B | | | | | | | | | | | | | | | |■| |
| 2.73620207438904 | R --> CDH1 | | |■| | | | | | | | | | | | | | |
| 2.74472737972518 | R --> CBFB | | | | | | | | |■| | | | | | | | |
| 2.75611055241523 | R --> RUNX1 | | | | | | |■| | | | | | | | | | |
| 2.75792533789356 | R --> MED23 | | | | | | | | | | | | | | |■| | |
| 2.78395266313118 | R --> TP53 | |■| | | | | | | | | | | | | | | |
| 2.79759691421067 | R --> ADAM29 | | | | | | | | | | |■| | | | | | |
| 2.79875481470716 | R --> MYO6 | | | | | | | | | | | | |■| | | | |
| 2.79941035546168 | R --> NPAS4 | | | | | | | | | | | | | |■| | | |
| 2.79943735321387 | R --> MAP3K1 | | | | |■| | | | | | | | | | | | |
| 2.8570705462381 | R --> GATA3 | | | |■| | | | | | | | | | | | | |
| 3.17292525307599 | R --> PIK3CA |■| | | | | | | | | | | | | | | | |
| 3.73787233597365 | R --> MAP2K4 --> ARID1A | | | | | |■| | | | | |■| | | | | |
| 3.83251698257823 | R --> CDH1 --> MAP2K4 | | |■| | |■| | | | | | | | | | | |
| 3.8613073440762 | R --> TBX3 --> CTCF | | | | | | | |■| |■| | | | | | | |
| 3.88461746616542 | R --> MAP2K4 --> MYO6 | | | | | |■| | | | | | |■| | | | |
| 3.8949899358325 | R --> TP53 --> MAP2K4 | |■| | | |■| | | | | | | | | | | |
| 3.89950276838655 | R --> TBX3 --> CBFB | | | | | | | |■|■| | | | | | | | |
| 3.90818772602653 | R --> CDH1 --> ARID1A | | |■| | | | | | | | |■| | | | | |
| 3.93472572846529 | R --> ARID1A --> MED23 | | | | | | | | | | | |■| | |■| | |
| 3.95195325645953 | R --> CDH1 --> TBX3 | | |■| | | | |■| | | | | | | | | |
| 3.98686025782634 | R --> TP53 --> TBX3 | |■| | | | | |■| | | | | | | | | |
| 3.99145712767097 | R --> GATA3 --> MAP2K4 | | | |■| |■| | | | | | | | | | | |
| 4.03746186808313 | R --> CTCF --> GRHL2 | | | | | | | | | |■| | | | | | |■|
| 4.10079284911008 | R --> MAP3K1 --> CTCF | | | | |■| | | | |■| | | | | | | |
| 4.10450246229066 | R --> CDH1 --> MED23 | | |■| | | | | | | | | | | |■| | |
| 4.10765251035681 | R --> TP53 --> CBFB | |■| | | | | | |■| | | | | | | | |
| 4.11090278934565 | R --> GATA3 --> TBX3 | | | |■| | | |■| | | | | | | | | |
| 4.11608659285386 | R --> CDH1 --> RUNX1 | | |■| | | |■| | | | | | | | | | |
| 4.11743135322005 | R --> CDH1 --> MAP3K1 | | |■| |■| | | | | | | | | | | | |
| 4.12323570184352 | R --> CDH1 --> MYO6 | | |■| | | | | | | | | |■| | | | |
| 4.12448020878165 | R --> TP53 --> RUNX1 | |■| | | | |■| | | | | | | | | | |
| 4.18895671615823 | R --> TP53 --> MAP3K1 | |■| | |■| | | | | | | | | | | | |
| 4.19608131076681 | R --> TP53 --> MYO6 | |■| | | | | | | | | | |■| | | | |
| 4.19876612289996 | R --> MAP3K1 --> MED23 | | | | |■| | | | | | | | | |■| | |
| 4.20123160438248 | R --> GATA3 --> CDH1 | | |■|■| | | | | | | | | | | | | |
| 4.20676411397217 | R --> TP53 --> ADAM29 | |■| | | | | | | | |■| | | | | | |
| 4.21850891562411 | R --> TP53 --> GRHL2 | |■| | | | | | | | | | | | | | |■|
| 4.22616158103902 | R --> MAP3K1 --> ADAM29 | | | | |■| | | | | |■| | | | | | |
| 4.27572848106676 | R --> GATA3 --> TP53 | |■| |■| | | | | | | | | | | | | |
| 4.30010585771875 | R --> GATA3 --> MAP3K1 | | | |■|■| | | | | | | | | | | | |
| 4.31074272813373 | R --> GATA3 --> CBFB | | | |■| | | | |■| | | | | | | | |
| 4.31312102189397 | R --> GATA3 --> NPAS4 | | | |■| | | | | | | | | |■| | | |
| 4.32200087691706 | R --> GATA3 --> MYO6 | | | |■| | | | | | | | |■| | | | |
| 4.48747504245117 | R --> PIK3CA --> TBX3 |■| | | | | | |■| | | | | | | | | |
| 4.50582697081201 | R --> PIK3CA --> ARID1A |■| | | | | | | | | | |■| | | | | |
| 4.53545610857787 | R --> PIK3CA --> MAP2K4 |■| | | | |■| | | | | | | | | | | |

| Time expectation | Order | PIK3CA | TP53 | CDH1 | GATA3 | MAP3K1 | MAP2K4 | RUNX1 | TBX3 | CBFB | CTCF | ADAM29 | ARID1A | MYO6 | NPAS4 | MED23 | ATP10B | GRHL2 |
|---|---|---|---|---|---|---|---|---|---|---|---|---|---|---|---|---|---|---|
| 4.54940027866825 | R --> PIK3CA --> CTCF |■| | | | | | | | |■| | | | | | | |
| 4.57754680164499 | R --> PIK3CA --> ATP10B |■| | | | | | | | | | | | | | |■| |
| 4.71396686038562 | R --> PIK3CA --> RUNX1 |■| | | | | |■| | | | | | | | | | |
| 4.76892041983634 | R --> PIK3CA --> TP53 |■|■| | | | | | | | | | | | | | | |
| 4.85595979132185 | R --> PIK3CA --> ADAM29 |■| | | | | | | | | |■| | | | | | |
| 4.8705837338389 | R --> PIK3CA --> MED23 |■| | | | | | | | | | | | | |■| | |
| 4.8872329151324 | R --> PIK3CA --> MAP3K1 |■| | | |■| | | | | | | | | | | | |
| 4.90908459903982 | R --> PIK3CA --> CDH1 |■| |■| | | | | | | | | | | | | | |
| 4.91115099833764 | R --> PIK3CA --> CBFB |■| | | | | | | |■| | | | | | | | |
| 4.9170501411738 | R --> PIK3CA --> GATA3 |■| | |■| | | | | | | | | | | | | |
| 4.99180690032654 | R --> PIK3CA --> GRHL2 |■| | | | | | | | | | | | | | | |■|
| 5.05923949345976 | R --> MAP3K1 --> TBX3 --> ARID1A | | | | |■| | |■| | | |■| | | | | |
| 5.12993755584062 | R --> GATA3 --> MAP2K4 --> TBX3 | | | |■| |■| |■| | | | | | | | | |
| 5.40835582689998 | R --> RUNX1 --> CTCF --> NPAS4 | | | | | | |■| | |■| | | |■| | | |
| 5.45135112279465 | R --> GATA3 --> ARID1A --> CBFB | | | |■| | | | |■| | |■| | | | | |
| 5.49267811247735 | R --> TP53 --> ATP10B --> ADAM29 | |■| | | | | | | | |■| | | | |■| |
| 5.50405826214877 | R --> GATA3 --> MAP3K1 --> TBX3 | | | |■|■| | |■| | | | | | | | | |
| 5.61865119942786 | R --> GATA3 --> CDH1 --> RUNX1 | | |■|■| | |■| | | | | | | | | | |
| 5.84243478713968 | R --> PIK3CA --> TBX3 --> CTCF |■| | | | | | |■| |■| | | | | | | |
| 6.0635279325792 | R --> PIK3CA --> TBX3 --> TP53 |■|■| | | | | |■| | | | | | | | | |
| 6.09836174362679 | R --> PIK3CA --> MAP2K4 --> ADAM29 |■| | | | |■| | | | |■| | | | | | |
| 6.10978482075964 | R --> PIK3CA --> MAP2K4 --> MED23 |■| | | | |■| | | | | | | | |■| | |
| 6.20476360637568 | R --> PIK3CA --> ARID1A --> GATA3 |■| | |■| | | | | | | |■| | | | | |
| 6.22058805007579 | R --> PIK3CA --> ARID1A --> CDH1 |■| |■| | | | | | | | |■| | | | | |
| 6.24032552364267 | R --> PIK3CA --> MAP2K4 --> CDH1 |■| |■| | |■| | | | | | | | | | | |
| 6.24747874181364 | R --> PIK3CA --> MAP2K4 --> GATA3 |■| | |■| |■| | | | | | | | | | | |
| 6.2968223360497 | R --> PIK3CA --> ATP10B --> CDH1 |■| |■| | | | | | | | | | | | |■| |
| 6.29720859035336 | R --> PIK3CA --> RUNX1 --> TP53 |■|■| | | | |■| | | | | | | | | | |
| 6.33827634923102 | R --> PIK3CA --> TBX3 --> CDH1 |■| |■| | | | |■| | | | | | | | | |
| 6.38980725968512 | R --> PIK3CA --> CTCF --> MAP3K1 |■| | | |■| | | | |■| | | | | | | |
| 6.4256617685557 | R --> PIK3CA --> NPAS4 --> TP53 |■|■| | | | | | | | | | | |■| | | |
| 6.45478392319306 | R --> PIK3CA --> RUNX1 --> MAP3K1 |■| | | |■| |■| | | | | | | | | | |
| 6.47771454706244 | R --> PIK3CA --> RUNX1 --> CBFB |■| | | | | |■| |■| | | | | | | | |
| 6.49314553818263 | R --> PIK3CA --> RUNX1 --> GATA3 |■| | |■| | |■| | | | | | | | | | |
| 6.50071207434945 | R --> PIK3CA --> MED23 --> TP53 |■|■| | | | | | | | | | | | |■| | |
| 6.5658566943657 | R --> PIK3CA --> TP53 --> CDH1 |■|■|■| | | | | | | | | | | | | | |
| 6.66035509702816 | R --> PIK3CA --> RUNX1 --> CDH1 |■| |■| | | |■| | | | | | | | | | |
| 6.68630442146548 | R --> PIK3CA --> MED23 --> MAP3K1 |■| | | |■| | | | | | | | | |■| | |
| 6.74336684967461 | R --> PIK3CA --> MAP3K1 --> CDH1 |■| |■| |■| | | | | | | | | | | | |
| 6.759665875532 | R --> PIK3CA --> CBFB --> CDH1 |■| |■| | | | | |■| | | | | | | | |
| 6.78084837500728 | R --> PIK3CA --> GATA3 --> CDH1 |■| |■|■| | | | | | | | | | | | | |
| 6.8344365039883 | R --> PIK3CA --> CBFB --> GRHL2 |■| | | | | | | |■| | | | | | | |■|
| 6.84292370358091 | R --> PIK3CA --> GATA3 --> CBFB |■| | |■| | | | |■| | | | | | | | |
| 7.38371891780409 | R --> PIK3CA --> ATP10B --> TP53 --> MAP2K4 |■|■| | | |■| | | | | | | | | |■| |
| 7.84656468747 | R --> PIK3CA --> MAP2K4 --> MED23 --> ADAM29 |■| | | | |■| | | | |■| | | |■| | |
| 8.00954329678745 | R --> PIK3CA --> ARID1A --> CDH1 --> MAP3K1 |■| |■| |■| | | | | | |■| | | | | |
| 8.15020411387736 | R --> PIK3CA --> GRHL2 --> CDH1 --> TBX3 |■| |■| | | | |■| | | | | | | | |■|
| 8.18953590180143 | R --> PIK3CA --> MED23 --> CDH1 --> TBX3 |■| |■| | | | |■| | | | | | |■| | |
| 8.53684038565906 | R --> PIK3CA --> GRHL2 --> CDH1 --> TP53 |■|■|■| | | | | | | | | | | | | |■|
| 9.18050534550109 | R --> PIK3CA --> ARID1A --> GATA3 --> MAP2K4 --> MYO6 |■| | |■| |■| | | | | |■|■| | | | |
| 9.34033833796437 | R --> PIK3CA --> ARID1A --> CDH1 --> ATP10B --> MYO6 |■| |■| | | | | | | | |■|■| | |■| |

Figure 1: Maximum likelihood estimation of the accumulation orders and waiting times for all unique profiles.

#### References


[1] Charles Van Loan. The sensitivity of the matrix exponential. *SIAM Journal on Numerical Analysis*, 14(6):971–981, 1977.

[2] Luca Dieci and Alessandra Papini. Padé approximation for the exponential of a block triangular matrix. *Linear Algebra and its Applications*, 308(1-3):183–202, 2000.

[3] Igor Najfeld and Timothy F Havel. Derivatives of the matrix exponential and their computation. *Advances in Applied Mathematics*, 16(3):321–375, 1995.
